# DNA epigenetic marks are linked to reproductive aberrations in amphipods

**DOI:** 10.1101/594788

**Authors:** Elena Gorokhova, Giulia Martella, Nisha H. Motwani, Natalia Y. Tretyakova, Brita Sundelin, Hitesh V. Motwani

## Abstract

Linking exposure to environmental contaminants with diseases is crucial for proposing preventive and regulatory actions. Upon exposure to anthropogenic chemicals, covalent modifications on the genome can drive developmental and reproductive disorders in wild populations, with subsequent effects on the population persistence. Hence, screening of chemical modifications on DNA can be used to provide information on the probability of such disorders in populations of concern. Using a high-resolution mass spectrometry methodology, we identified DNA nucleoside adducts in gravid females of the Baltic amphipods *Monoporeia affinis*, and linked the adduct profiles to the frequency of embryo malformations in the broods. Twenty-three putative nucleoside adducts were detected in the females and their embryos, and eight modifications were structurally identified using high-resolution accurate mass data. To identify which adducts were significantly associated with embryo malformations, partial least squares regression (PLSR) modelling was applied. The PLSR model yielded three adducts as the key predictors: methylation at two different positions of the DNA (5-methyl-2’-deoxycytidine and N^6^-methyl-2’-deoxyadenosine) representing epigenetic marks, and a structurally unidentified nucleoside adduct. These adducts predicted the elevated frequency of the malformations with a high classification accuracy (84%). To the best of our knowledge, this is the first application of DNA adductomics for identification of contaminant-induced malformations in field-collected animals. The method can be adapted for a broad range of species and evolve as a new omics tool in environmental health assessment.

## INTRODUCTION

Covalent DNA adducts can be formed by chemicals that are electrophilic as such or those that form reactive metabolites^1-3^. Another pathway for modification of the DNA is through oxidation by reactive oxygen species (ROS) formed under oxidative stress^4,5^. If not repaired, such DNA lesions can interfere with the accuracy of DNA polymerases during replication which could lead to mutations. Alternatively, environmental exposures can dysregulate epigenetic modifications of DNA bases, such as 5-methylcytosine. Unlike the genotoxic DNA adducts, epigenetic DNA modifications are introduced enzymatically and are involved in regulating the levels of gene expression^6-8^. Such genetic and epigenetic modifications can potentially lead to deleterious growth and developmental aberrations in wild populations, which may, in turn, result in reproductive failure and population instability. However, studies linking genotoxic exposures to reproductive disorders in environmental samples are rare^9,10^, and little is known about the relationships between specific types of DNA modifications and organism-level responses.

A comprehensive analysis of the modifications on the nucleobase moieties of DNA is addressed by DNA adductomics, an emerging field in systems toxicology. High-resolution mass spectrometry (HRMS) methods employing Orbitrap MS instrumentation have gained popularity in the adductomics analysis due to their ability to detect adducts with high mass accuracy in complex mixtures^11,12^. However, DNA adductomics being a relatively young field presents many analytical challenges, including optimization of chromatographic separations, structural assignments for novel adducts, and data interpretation.

Human activities over the past decades have aggravated the environmental pollution in many estuarine systems, including the Baltic Sea^13^. Physical adsorption causes organic and inorganic contaminants to become highly enriched in sediments and porewater as compared to the water column^14^. This makes the benthic animals, such as amphipods, polychaetes, and clams, particularly vulnerable to continuous exposure to the sediment-bound compounds. Such organisms are thus useful as sentinels in environmental monitoring programs.

The amphipod *Monoporeia affinis* is a keystone species in soft bottoms of the Baltic Sea, where it often dominates benthic communities. Their great abundance, high quality as prey, and active feeding on organic material in the sediments make these animals an important trophic link in the transfer of sediment-associated contaminants to the benthic and pelagic food webs. In this semelparous amphipod, contaminant exposure can induce various reproductive disorders, including embryonic aberrations^15^. Moreover, several aberration types were linked to specific contaminants in the ambient sediments, providing a basis for application of the embryo aberration analysis in the environmental assessment^16,17^. Currently, embryonic aberrations in the amphipods are used as an indicator for biological effects of contaminants within the Swedish National Marine Monitoring Program (SNMMP) and as a supporting indicator for Descriptor 8 (Hazardous substances) in the Marine Strategy Framework Directive (MSFD).

Towards the development of the screening methods that would facilitate detecting biological effects of environmental contaminants, we applied HRMS-based DNA adductomics in *M. affinis* collected in the northern Baltic Sea. We hypothesized that reproductive aberrations in these amphipods are a manifestation of toxicity related to specific DNA modifications. To test this hypothesis, we evaluated whether certain modifications of the DNA were associated with high frequency of reproductive aberrations.

## METHODS

### Chemicals and other materials

Deoxyribonucleic acid from calf thymus (ctDNA) sodium salt, 2’-deoxyguanosine (dG), 2’-deoxycytidine (dC), 2’-deoxyadenosine (dA), thymidine (T), 5-methyl-2’-deoxycytidine (5-me-dC), 8-oxo-7,8-dihydro-2’-deoxyguanosine (8-oxod-G), N^6^-methyl-2’-deoxyadenosine (N^6^-me-dA), nuclease P1 from *Penicillium citrinum* (NP1), phosphodiesterase I from *Crotalus adamanteus* (snake) venom (SVPDE), alkaline phosphatase from *Escherichia coli* (AKP), ammonium acetate, ammonium bicarbonate, 2,2,6,6-tetramethyilpiperidine-1-oxyl (Tris-buffer), zinc chloride and formic acid were obtained from Sigma-Aldrich (St. Louis, MO). Chelex-100 resin was purchased from Bio-Rad (Solna, Sweden). All solvents used were of HPLC grade. Experiments concerning DNA were carried out in DNA LoBind tubes, 1.5 mL (Eppendorf).

### Amphipod samples

As a part of SNMMP, gravid *M. affinis* were collected in the middle of January 2017 at 30 stations along the Swedish coast, from the Quark in the north of the Bothnian Sea to Western Gotland Basin (Supplementary information, **Fig. S1**). These amphipods reproduce in late-November/early-December and carry their embryos in a brood pouch (marsupium) until offspring release in February. In the end of January, the embryos reach advanced developmental stage of 4–8, making it possible to identify developmental aberrations. Gravid females were collected by a benthic sled^18^, sorted, and transported in ambient water (<7 □) to the laboratory for embryo analysis.

### Embryo aberration analysis

Eggs and embryos were removed from the marsupium of the female, and embryos were examined using a stereo-microscope (Leica M10 80× magnification) with polarized cold light. For each female, we recorded fecundity (eggs per female), embryo developmental stage, and number of aberrant embryos, if present. The aberration types considered were all incompatible with survival^17^. For both females and their embryos, two groups were created: (1) healthy (aberration frequency ≤ 5%; 21 females and their broods), and (2) unhealthy (aberration frequency 8-41%; 19 females and their broods). To assign the amphipods to these groups, the threshold for aberration frequency of 6%, i.e., percentage of malformed embryos of the total number of embryos in the brood, was applied based on the evaluation of the background frequency of embryo aberrations in this species^19^.

In the DNA analysis, the females with the brood removed and their embryos were treated separately. Each female was snap frozen in liquid nitrogen directly after the brood removal, and the dissected broods were treated in the same way directly after the embryo analysis. The samples were stored at −80 °C until the DNA extraction.

### DNA extraction

Individual samples were manually homogenized using a Kontec pestle, and genomic DNA was extracted with 350 μL of 6% Chelex-100 solution^20^. Briefly, the homogenate and chelex solution were incubated at 65 °C for 3 h and intermittently vortexed. After centrifugation at 14000 rpm for 10 min, the supernatant was transferred to an Eppendorf tube and stored at 4 °C overnight. DNA concentrations and purity were determined with a Nanophotometer™ (Implen), which gave A260/A280 as 1.7–1.8. The extracted DNA was then stored at −20 °C until sample preparation for liquid chromatography mass spectrometry (LC-MS) analysis.

### Enzymatic digestion

Amphipod DNA (15 µg) extracted from the individual females and their broods was mixed with Tris-buffer (1 mM, pH 7.4) to give a total volume of 300 µL. Ammonium acetate (0.1 M, 30 µL), zinc chloride (10 mM, 12 µL) and ammonium bicarbonate (1 M, 35 µL) were added along with the enzymes for digestion, which were NP1 (0.1 U/µL, 8 µL), SVPDE (0.000126 U/µL, 7 µL) and AKP (0.029 U/µL, 7 µL). The mixture was incubated at 37 °C for 2.5 h, followed by centrifugation at 14000 rpm (4 °C, 10min). The supernatant containing the nucleosides was transferred to a septum sealed vial and analyzed by LC-MS. Further, blank samples were obtained with the procedure being repeated in triplicates but without any DNA.

### Liquid chromatography conditions

The LC-MS system used consisted of a Dionex UltiMate 3000 LC device interfaced to an Orbitrap Q Exactive HF HRMS (Thermo Fisher Scientific, MA). The mobile phase for high-pressure liquid chromatography (HPLC) consisted of a mixture of water-methanol; system A with 5% methanol and system B with 95% methanol, each containing 0.1% formic acid. The HPLC column used was a Superlco Ascentis® Express F5 2.7 Micron HPLC column (15cm x 2.1mm) from Sigma-Aldrich. Sample injection volume was 20 μL. A flow rate of 120 μL/min was employed with the column temperature maintained at 25 °C. The LC gradient consisted of an initial 2 min equilibration at 5% of B, which was increased to 30% in 6 min and then to 100% in 2 min. After holding at this composition for 2 min, a ramping to the initial condition of 5% B was done in 1 min, and the system was re-equilibrated for 3 min before the next injection. An automated switch-valve was connected between the LC column and the Orbitrap MS, which was set to allow the eluent from the column to enter the waste during the first minute after injection to remove polar impurities in the samples. After this one minute, the valve was used to divert the eluent into the ion source of the mass spectrometer.

### Orbitrap HRMS/MS analysis

HRMS analysis was carried out using an Orbitrap Q Exactive HF mass spectrometer equipped with a heated electrospray ionization (HESI) source. The HESI MS parameters were initially tuned for maximal signal intensity by infusion using a standard mixture prepared that contained dG, dA, dC and T (10 fmol/µL of each). The optimized MS parameters that were used for all further experiments were as follows. Spray voltage, 3.5 kV; spray current, 22 µA; capillary temperature, 275 °C; sheath gas, 20 arbitrary units (au); auxiliary gas, 10 au; S-Lens RF level, 60%; and probe heater temperature, 240 °C. The mass spectrometer was operated in the positive ion mode using normalized collision energy of 30 eV. Screening for the DNA modifications as nucleoside adducts was performed using full scan (FS) / data-independent acquisition (DIA) mode, wherein MS/MS fragmentation is performed on a sequential scan range. The full MS scanning was conducted with a mass resolution of 120000, automatic gain control (AGC) target 3e^6^, maximum ion injection time (IT) 200 ms, and scan range from 110 to 1200 m/z. The DIA was set to have resolution 60000, AGC target 5e^5^, maximum ion IT 120 ms, loop count 31, and scan range from 195 to 355 m/z, which was divided into 16 discrete m/z ranges with an isolation window of 10 m/z, in the form 200 ± 5, 210 ± 5, and up to 350 ± 5 m/z.

### Data processing

The generated data was processed using XCalibur 3.1 (Thermo Fisher Scientific). Each mass range from the DIA mode was manually evaluated in the Qual browser by monitoring for the fragment ion with *m/z* 117.0552 ± 5ppm, corresponding to the protonated deoxyribose ion, [dR]^+^, to select the adduct candidates. The candidates were filtered as possible nucleoside adducts when two others ions, in addition to the [dR]^+^, were detected in the form of parent ion [M]^+^ and a specific fragment ion [M-dR]^+^ (**Fig. S2**) within the same chromatographic peak, for which the following equation sustained within a 5ppm mass tolerance for each ion; [M]^+^ = [M-dR]^+^ + 116.0473. The corresponding FS spectra were used for confirmation of the parent ions. This processing was initially applied to generate a list of putative adducts from a set of representative samples, five females and five embryos from each group of healthy and unhealthy. Quantification was then performed in XCalibur’s Quan browser for the putative adducts (A1□A23) in all samples. Peak areas were measured using extracted ion chromatograms (EIC) of the adducts corresponding to m/z (± 5ppm) of the specific fragments which had higher abundance than the parent ions under the employed MS conditions. A normalized maximum intensity level (NL) of 1.0E3 was set as an apparent limit of quantification for individual peak areas. All adducts below this limit (A5 and A10 in females; A1, A5, A10, and A22 in embryo broods) were excluded from further statistical analysis. The measured peak areas were normalized to that of dG (adduct area × 10^2^ / dG area) from the same sample, followed by statistical analysis. Normalization to dG, which has also been successfully applied in other LC-MS based studies including those investigating changes in 5-me-dC levels^21,22^, circumvents for loses during sample preparation and avoids the need for expensive isotope-labeled internal standards.

### Identification of DNA modifications

Preliminary identification of each nucleoside adduct was based on the elemental composition obtained from the high mass accuracy data of the parent ions, and their comparison with the corresponding theoretical values calculated from ChemDraw Professional 16.0. The identification included calculating for protonated and sodiated adducts. In addition to the four unmodified nucleosides, the identities of 5-me-dC, N^6^-me-dA, and 8-oxo-dG were confirmed using authentic reference standards. Standard solutions of each reference compound (100 fmol/µL) were prepared in deionized water and analyzed by LC-Orbitrap employing the same settings as described above. The resulting HRMS data, as well as retention time on the chromatogram, were compared with the corresponding data obtained from the amphipod samples. Spiking of amphipod samples with respective standard solution was performed to further confirm the identification of 5-me-dC, N^6^-me-dA, and 8-oxo-dG.

### Evaluation of DIA method for quantification

Adduct measurements based on the DIA method was evaluated for repeatability and variability using ctDNA, a commercially available DNA standard. Five replicates of ctDNA (15 µg) were dissolved in Tris-buffer (1mM, pH 7.4, 300 µL) and enzymatically digested in the same way as the amphipod DNA. The resulting mixture was analyzed by LC-Orbitrap using the same DIA settings as described above. Peak areas were measured using EIC corresponding to the specific fragments, and normalized to dG, for five nucleoside adducts covering the selected mass range and including the identified adducts. These included parent ions having m/z 242.1146 (5-me-dC), 284.0983 (8-oxo-dG), 289.1758 (A7), 300.1299 (A12), 306.0603 (A17); N^6^-me-dA was below quantification limit in the ctDNA samples and hence not selected in the evaluation. The standard deviation (SD) and coefficient of variation (CV) were calculated for each measured analyte from the ctDNA (**Fig. S3**).

### Statistical analysis

The full dataset consisted of the putative nucleoside adduct profile in the female and their embryos and the health status as a group variable. Adducts A4, A13, A16, and A18 were excluded from the modelling, since these were attributed as sodiated adducts and had high correlation with their corresponding protonated forms. To evaluate differences in frequencies of specific adducts between the females and embryos and between the healthy and unhealthy individuals, we applied the following statistical methods. To compare the differences in the profiles of adducts between the females and their embryos, the data variability across the samples was visualized using non-dimensional multiple scaling (NMDS) based on Bray–Curtis distances. Permutational Multivariate Analysis of Variance using distance matrices, PERMANOVA, with 1000 Monte Carlo permutations was performed based on Bray–Curtis distances^23^ and using software PRIMER+ v.6 (Primer-e, Ivybridge, UK). Further, to produce a predictive model for the amphipod reproductive health using adducts, we combined projection to latent structures using partial least squares (PLS-DA) and a follow-up logistic regression. Whereas the standard logistic regression model has difficulty handling a large number of variables, PLS-DA can be constructed using all adducts, without any prior variable selection step. The standard logistic regression model is then constructed using the selected adducts identified by PLS-DA^24^. Therefore, PLS-DA was applied to search for discriminative adducts that contributed to the separation between the adduct profiles (DNA modifications across an individual) belonging to different groups (healthy vs. unhealthy) as implemented in MetaboAnalyst^25^. The influential predictors were selected based on their VIP values with a cutoff of 0.8. The predictors were then used in the logistic model with the health status as a binomial response variable, binomial error structure, log-link function, and the AIC score as the selection criterion. Odds ratios of the logistic regression were used as a measure of associations between the treatment and the predictors. The sensitivity (i.e., true positives, or the proportion of cases correctly identified by the model as being unaffected by stress) and the specificity (e.g., true negatives, the proportion of cases correctly identified by the model as experiencing stress) were extracted from the ROC curves.

## RESULTS

### Detection of DNA modifications using high-resolution mass spectrometry

To screen for potential DNA modifications, released as nucleoside adducts in the enzymatic digests, we took advantage of the characteristic dR moiety in the LC-HRMS analysis on an Orbitrap MS system. Twenty-three putative nucleoside adducts (abbreviated as A1 to A23; **Table 1**) were detected in the amphipods. The detection was based on fragmentation of the nucleoside adducts, using electrospray ionization in positive ion mode, to m/z 117.0552 (corresponding to protonated dR) and a specific fragment for each nucleoside adduct corresponding to nucleobases with structural modifications at the purine or pyrimidine ring (**Fig. S2**). High mass accuracy within a 5ppm range was employed, facilitating structural identification of the unknown adducts based on their elemental composition.

**Table 1.** Characterization of the detected nucleoside adducts, and the 4 unmodified nucleosides, using a HRAM adductomics approach. Mass-to-charge ratio of observed molecular ion [M]^+^ and that of the fragment [M-dR]^+^, and retention time under the employed chromatographic conditions are given. For the identified compounds, the mass difference between that observed (reported in this table) and calculated m/z was less than 3ppm.

| Detected nucleosides | Molecular ion, $m/z [M]^+$ | Specific fragment, $m/z [M-dR]^+$ | Retention time, min | Identified compounds |
| --- | --- | --- | --- | --- |
| A1 | 295.1203 | 179.0739 | 3.51 |  |
| A2 | 278.1605 | 162.1126 | 3.59 |  |
| dC | 228.0978 | 112.0507 | 3.81 | dC<sup>a</sup> |
| A3 | 242.1146 | 126.0664 | 3.97 | 5-me-dC<sup>a</sup> |
| A4 | 264.0961 | 148.0484 | 3.98 | Na adduct of 5-me-dC<sup>b</sup> |
| A5 | 244.0932 | 128.0457 | 4.48 | 5-OH-dC |
| A6 | 229.0818 | 113.0348 | 5.49 | dU |
| A7 | 289.1758 | 173.1288 | 5.60 |  |
| A8 | 268.1043 | 152.0569 | 6.40 | 8-OH-dA |
| dA | 252.1094 | 136.0620 | 6.41 | dA<sup>a</sup> |
| A9 | 239.1139 | 123.0667 | 6.41 |  |
| A10 | 274.0915 | 158.0440 | 6.42 | Na adduct of dA<sup>b</sup> |
| A11 | 284.1358 | 168.0883 | 6.43 |  |
| A12 | 300.1299 | 184.0832 | 6.43 |  |
| A13 | 275.0753 | 159.0280 | 7.25 | Na adduct of dI |
| A14 | 253.0931 | 137.0460 | 7.33 | dI |
| dG | 268.1033 | 152.0568 | 7.51 | dG<sup>a</sup> |
| A15 | 274.1148 | 158.0676 | 7.51 | Gh |
| A16 | 290.0852 | 174.0389 | 7.52 | Na adduct of dG<sup>b</sup> |
| A17 | 306.0603 | 190.0129 | 7.61 |  |
| T | 243.0970 | 127.0504 | 7.99 | T<sup>a</sup> |
| A18 | 265.0798 | 149.0324 | 8.00 | Na adduct of T<sup>b</sup> |
| A19 | 327.0504 | 211.0029 | 8.03 |  |
| A20 | 236.1281 | 120.0809 | 8.48 |  |
| A21 | 284.0989 | 168.0518 | 8.75 | 8-oxo-dG<sup>a</sup> |
| A22 | 266.1257 | 150.0777 | 9.42 | N<sup>6</sup>-me-dA<sup>a</sup> |
| A23 | 252.1228 | 136.0759 | 11.34 |  |

<sup>a</sup>Identification confirmed by comparison with respective standards.
<sup>b</sup>These were assigned as sodiated adducts of the respective compounds (based on retention time and molecular masses), with molecular ion and specific fragment represented as $[M+Na]^+$ and $[M-dR+Na]^+$ , respectively. All others are assigned as protonated adducts with molecular ion and specific fragment represented as $[M+H]^+$ and $[M-dR+H]^+$ , respectively.

The amount of digested DNA injected on-column was kept constant (0.75 µg), which was judged to be sufficient for adduct detection. No nucleosides were detected in the blank samples prepared by the same procedure but in the absence of DNA. Representative chromatograms corresponding to adducts detection in amphipod samples using LC-HRMS/MS analysis are shown in **Fig. 1**. The figure also highlights the advantage of using HRMS for resolving nucleoside adducts from the complex sets of ions present in the samples (**Fig. 1 C**), as high background from sample matrix was observed when using low mass resolution settings (**Fig. 1 B**).

**Figure 1.**
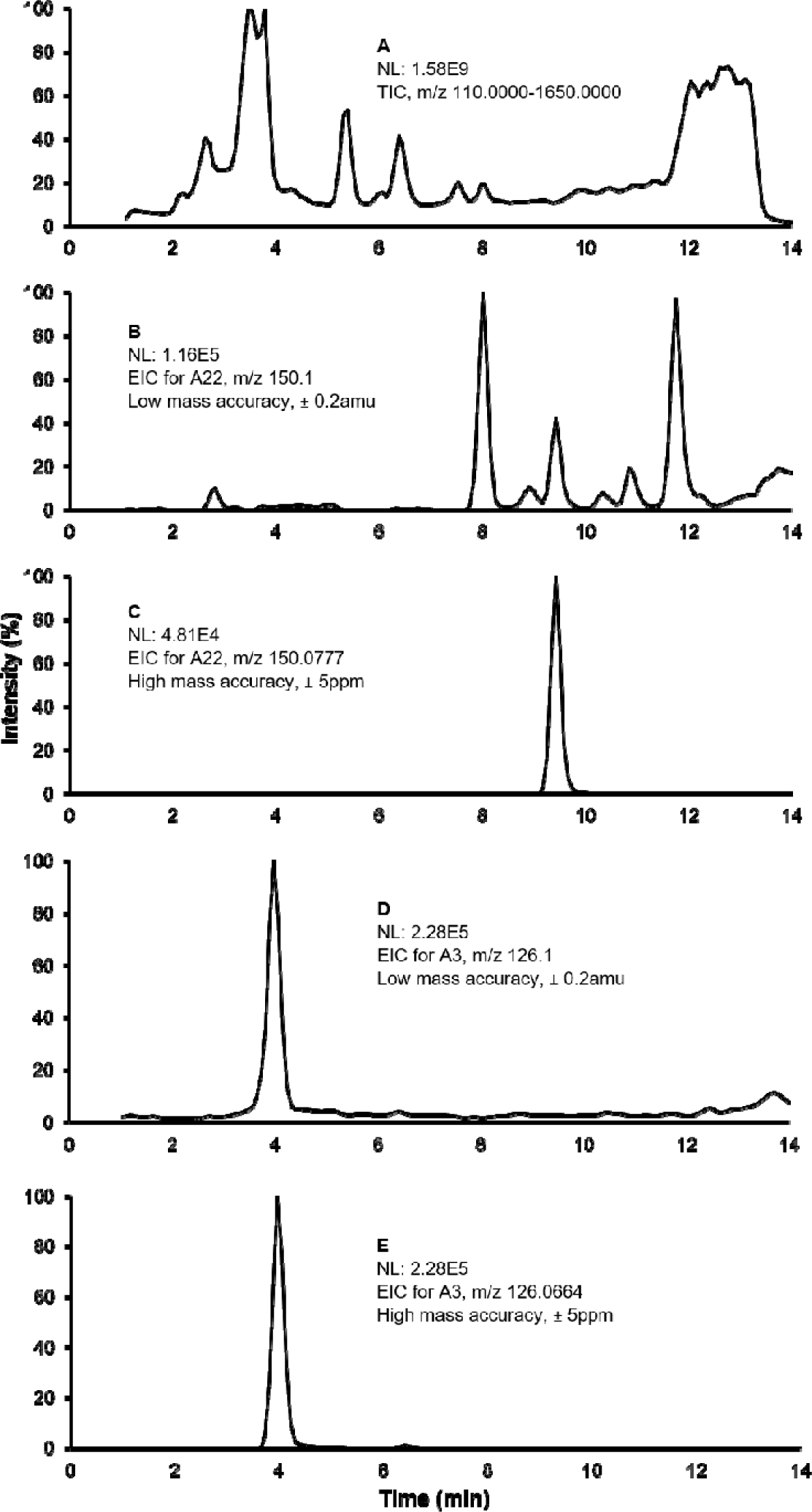
Representative chromatograms from an amphipod showing differences in high and low mass accuracy for resolving of nucleoside adducts. **A**: Total ion chromatogram (TIC) of the full scan data showing high background signal from the sample matrix, which does not allow for detection of the adducts; NL, normalized maximum ion intensity level. **B**: Extracted ion chromatogram (EIC) corresponding to [M-dR+H]^+^ for A22 (N^6^-me-dA) at a mass tolerance typical of quadrupole instrument (± 0.2 amu). The peak corresponding to A22 is not clearly distinguishable. **C**: EIC corresponding to [M-dR+H]^+^ for A22 at a mass tolerance of 5ppm. The peak corresponding to the adduct is clearly resolved. **D** and **E**: Similar to B and C, respectively, but for A3 (5-me-dC). The peak is resolved in both panes.

### Structural identification of DNA modifications

Structural characterization of each nucleoside adduct was based on molecular weight information, characteristic fragment ions containing modified free base (**Fig. S2**), and HPLC retention time, which were compared to those of authentic standards (**Table 1)**. High resolution accurate mass (HRAM) measurements on an Orbitrap MS allowed for determination of possible elemental composition of the unknown adducts. Out of the 23 detected nucleoside adducts, 8 were structurally identified. **Fig. 2** shows the structures of nucleoside adducts that were identified in the amphipod DNA, along with their elemental composition and calculated masses. The mass difference between data recorded for the standards (5-me-dC, N^6^-me-dA, and 8-oxo-dG) and the corresponding analytes from amphipod samples was < 1ppm. When amphipod DNA digested samples were spiked with standard solutions of 5-me-dC, N^6^-me-dA, and 8-oxo-dG, complete co-elution of the analyte peaks was observed before and after spiking, confirming their identification in amphipod DNA (**Fig. S4**).

**Figure 2.**
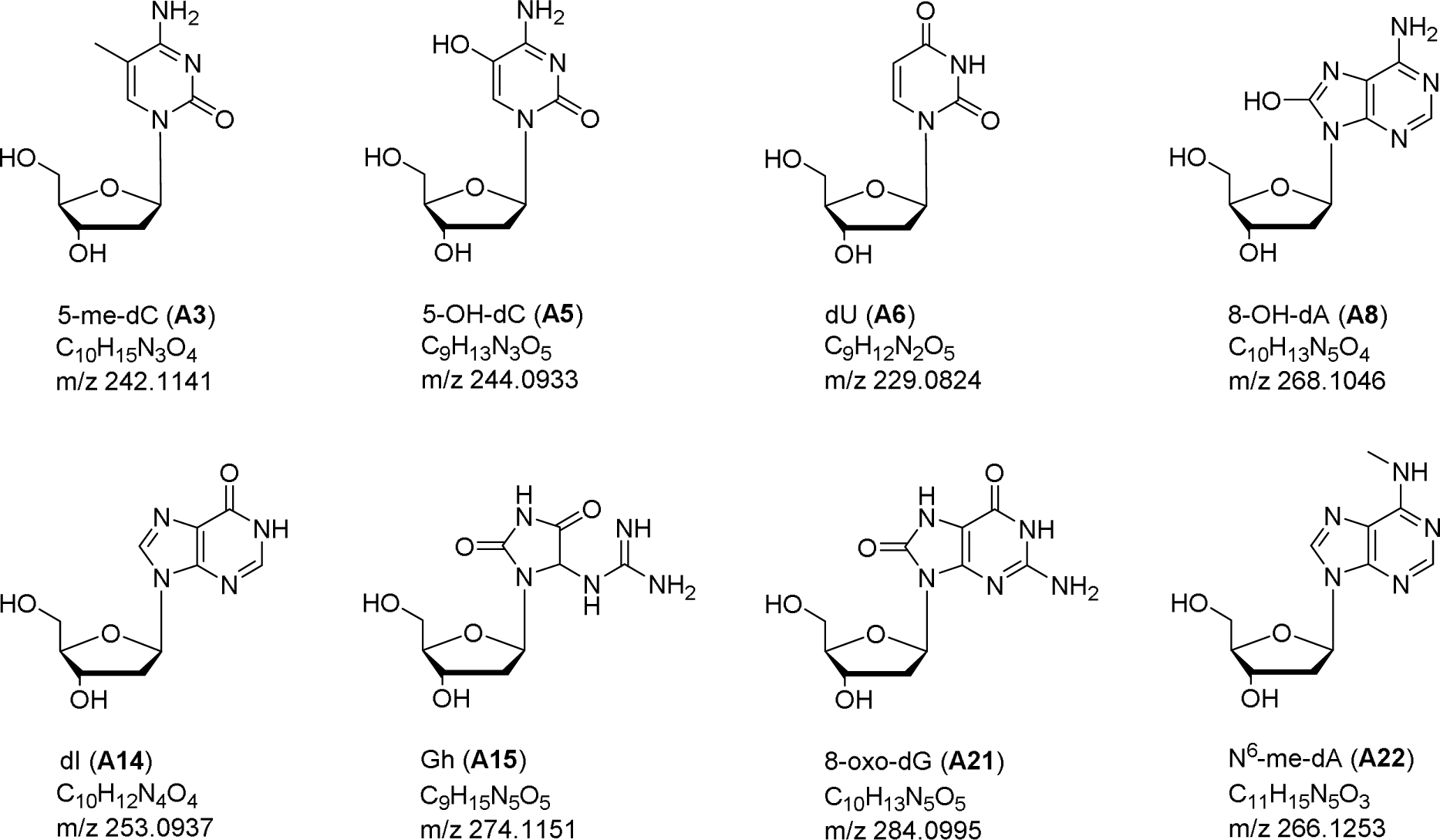
Chemical structure, elemental composition and calculated m/z [M+H]^+^ of nucleoside adducts identified in the amphipods. The identification of A3, A21 and A22 were confirmed by comparison with respective standards; the proposed structures of the others are based on the HRAM data.

### Quantitative evaluation of DNA modifications in females and embryos

For the majority of nucleoside adducts, their composition and abundance were similar between the females and the embryos (**Fig. S5 and S6**). The frequent adducts were distributed similarly across the samples, with A11 and A20 being amongst the most commonly detected nucleoside adducts in both females and embryos (**Fig. 3**). However, differences in the detection frequencies for some specific adducts between the females and embryos were also found. The NMDS plot of the Bray– Curtis distances confirmed that samples clustered primarily in a source-dependent manner along the NMDS2 axis, with variability between the broods (embryo samples) being significantly higher compared to that between the individual females (**Fig. 4**). Both NMDS stress value and ANOSIM output indicated significant differences in the nucleoside adduct profile between the females and their embryos (**Fig. 4**). Moreover, significant correlations were observed between different adduct types, such as A7, A20, and A23 (**Fig. S7**). Also, the detection frequencies correlated significantly positively between the females and their embryos for A15, A20, and A23 (**Table S1**).

**Figure 3.**
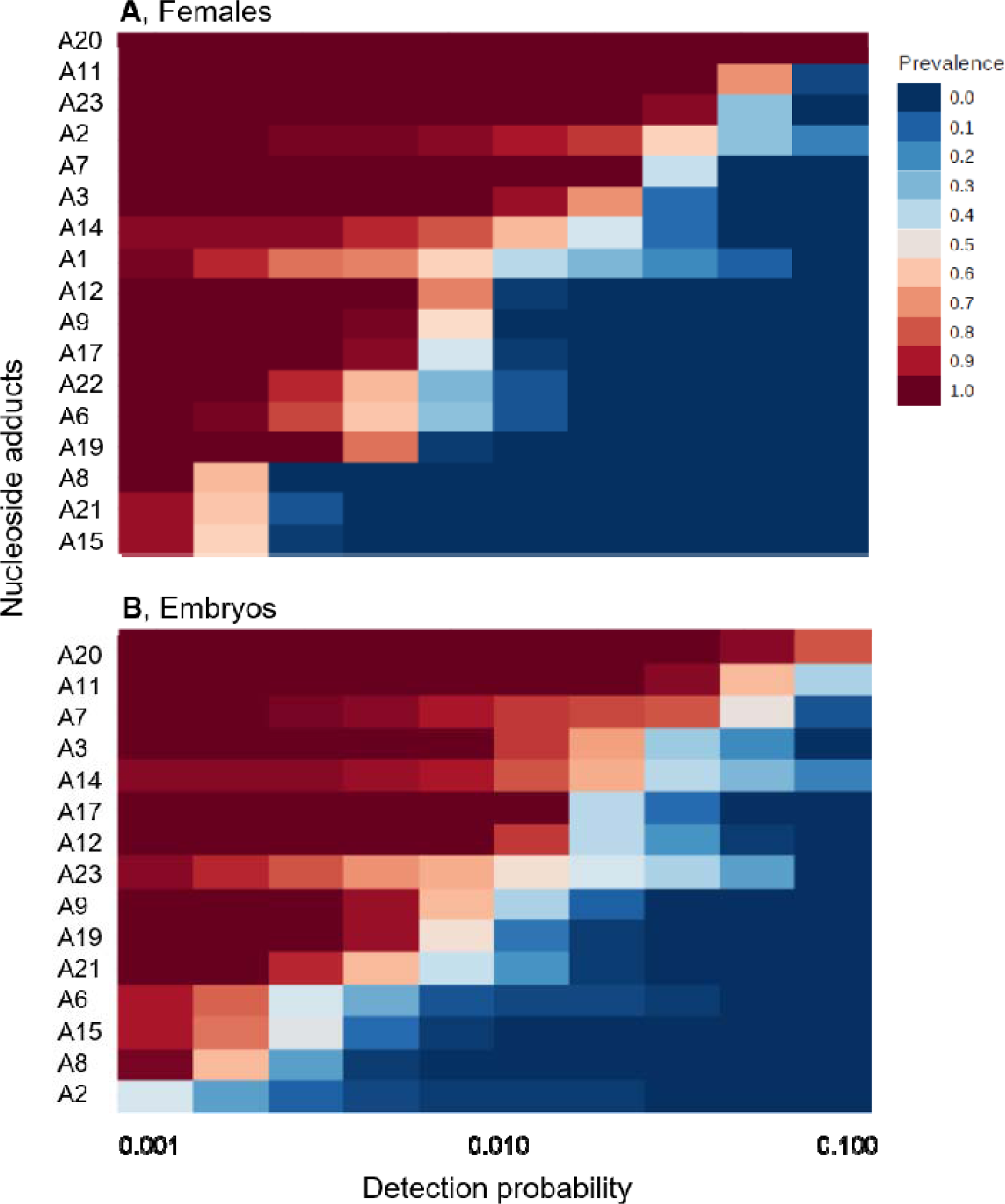
Profiles of nucleoside adducts in the females (**A**) and their embryos (**B**) expressed as their detection probability in 40 individual samples. See **Table 1** for additional information on specific adducts.

**Figure 4.**
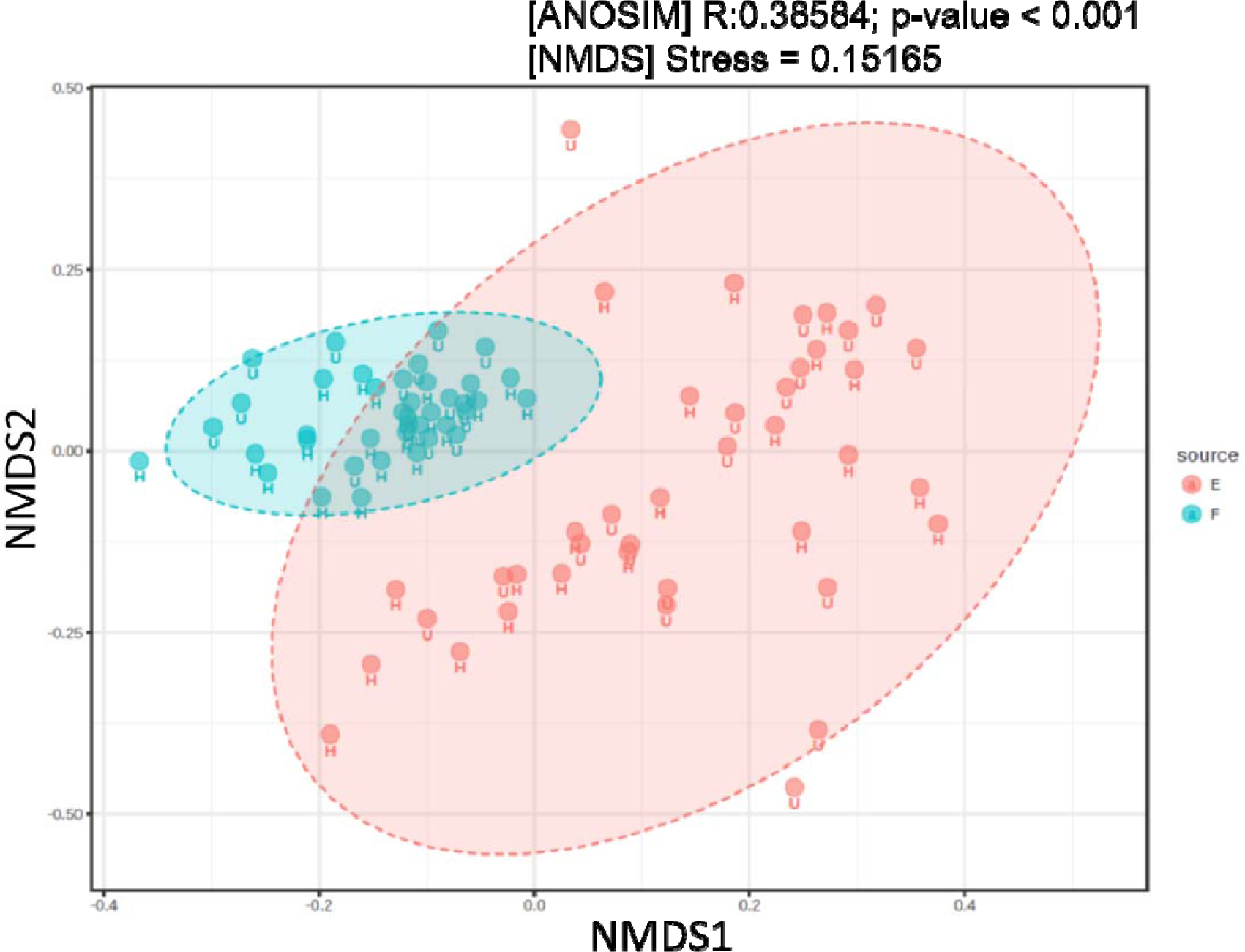
NMDS ordination diagram based on the nucleoside adducts in the females and their embryos. PERMANOVA results on the amount of variation in the adduct data attributable to the source (females vs. embryos) are based on Bray-Curtis dissimilarity index.

### DNA modifications as predictors of amphipod health status

Partial least squares-discriminant analysis (PLS-DA) using normalized peak areas for nucleoside adducts as predictors and amphipod health status (healthy vs. unhealthy) discriminated between the females carrying high number of malformed embryos (8-40%; unhealthy) and those with the background levels of malformations (≤ 5%; healthy) (**Fig. 5A**); the between-group difference was validated by the permutation test (p < 0.013; **Fig. 5B**). Three component matrices accounted for 32.1, 15.2, and 5.9 % of the total variance (**Fig. 5A**). Top contributors to the PLS-DA included A22 N^6^-me-dA), A9, A3 (5-me-dC), A12, and A1 in the females and A15 (Gh) and A7 in the embryos. Individually, among the female-associated adducts, A22, A9, and A3 were the most elevated, while A12 and A1 were the most depleted adducts in unhealthy individuals. Among the embryo-associated adducts, A7 was elevated while A15 was depleted in the unhealthy individuals (**Fig. 5C**). These adducts were the greatest contributors to the PLS-DA components 1–3 based on variable importance in projection (VIP) (**Fig. 5C**).

**Figure 5.**
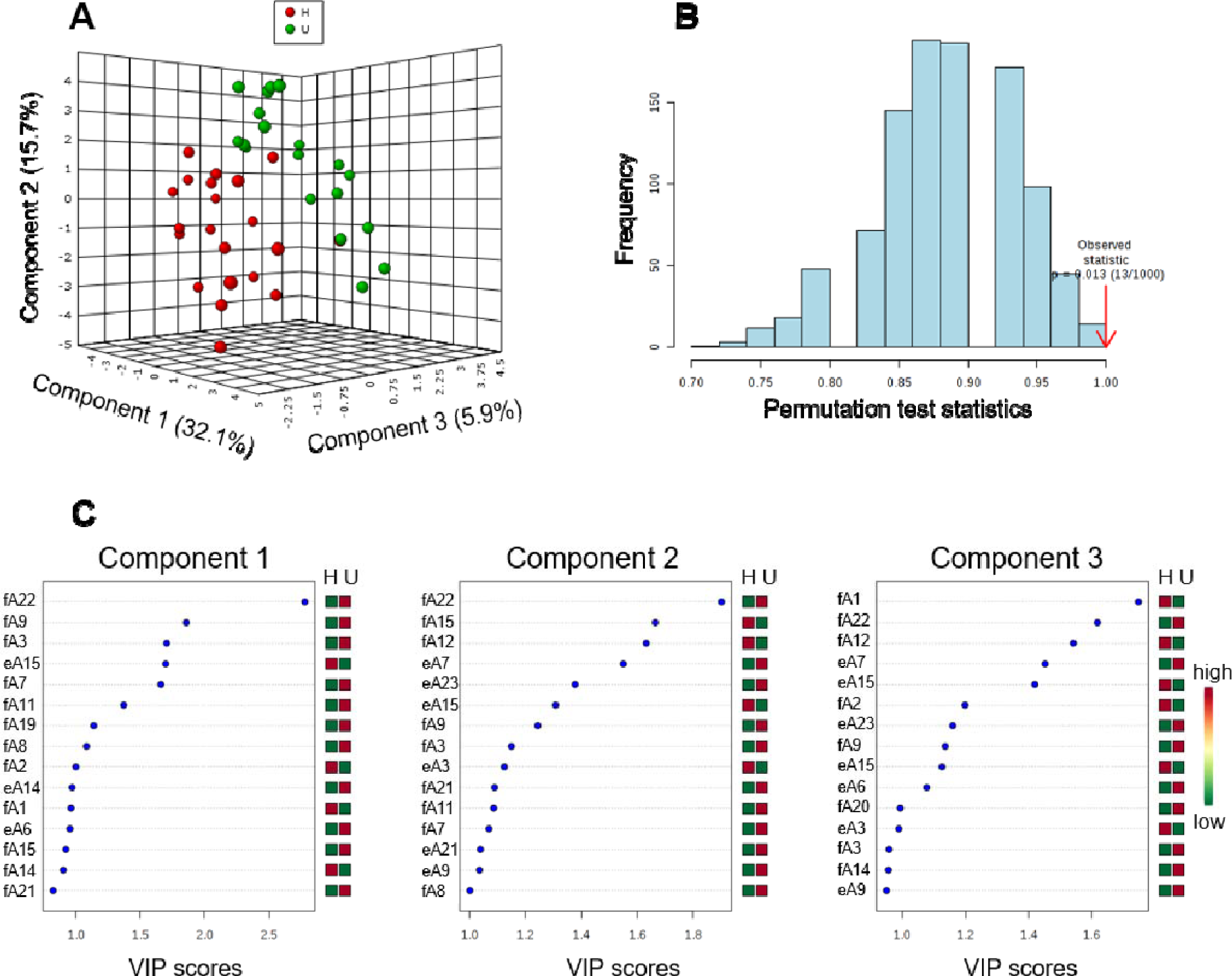
Partial least squares-discriminant analysis (PLS-DA) of nucleoside adducts in healthy (≤ 5% embryo aberrations) and unhealthy (8-41%) females. **A**: 3-Dimensional score plot of PLS-DA using components 1, 2, and 3, accounting for 32.1, 15.7, and 5.9 % of the total variance; **B**: Validation of PLS-DA by permutation test (p < 0.013); and **C**: Variable importance in projection (VIP) scores of 15 top contributors to PLS-DA, components 1–3; note the differences in the scale of the x-axis between the panels. The adducts (**Table 1**) measured in females and embryos are denoted as fA# and eA#, respectively.

A logistic model discriminated between healthy and unhealthy females with high classification accuracy (**Table S2**). Using this approach, the area under the receiver operating characteristic (ROC) curve (AUC) identified A22 (N^6^-me-dA), A9, and A3 (5-me-dC) to have the greatest specificity and sensitivity for distinguishing between reproductively unhealthy and healthy females based on their adductome (**Fig. S8**). For all the three adducts, increased levels were associated with high frequency of reproductive aberrations. In contrast, none of the adducts in the embryos that were identified as significant predictors for PLS-DA were included in the winning logistic model.

## DISCUSSION

Despite the biological plausibility of a link between exposure to environmental contaminants, reproductive abnormalities, and DNA modifications, our knowledge of these causal linkages is sparse, particularly for non-model species and in wild populations. The evidence is accumulating that environmental chemicals that exhibit reproductive toxicity do so at least in part through genotoxic mechanisms and epigenetic alterations in genomic DNA^9,26^. Here, we tested the hypothesis that the frequency of reproductive aberrations in Baltic Sea amphipods is related to the levels of certain DNA modifications in females and their embryos. To screen for DNA modifications, as nucleoside adducts, we employed an untargeted adductomics approach using HRMS, and the resulting data with high mass accuracy was used for identification of the adducts.

We found significant differences in the nucleoside adduct profile between the amphipods with high and low frequencies of the abnormal embryos in the brood and identified the adducts contributing most to these differences. Thus, the hypothesized relationship between the reproductive pathologies in the amphipods and the occurrence of DNA modifications was supported. The occurrence of the aberrant embryos was positively associated with relatively increased levels for 5-me-dC (A3) and N^6^-me-dA (A22), the epigenetic DNA modifications that are introduced enzymatically and have a regulatory role in cell functioning. We suggest that environmental exposure lead to epigenetic deregulation, which in turn caused developmental and reproductive toxicity in the animals. To the best of our knowledge, this is the first study demonstrating the utility of HRAM for DNA adductomics in wild populations. The method is applicable for hazard assessment of environmental contaminants and biological effect monitoring.

LC-MS/MS has become the preferred method for measuring DNA adducts due to the high selectivity obtained in comparison to, e.g., ^32^P-post-labeling technique^27-29^. Moreover, recent advances in MS instrumentation providing HRAM data have resulted in gaining more information on the chemical structure of the adduct based on its fragmentation pattern and more accurate elemental composition determination^12,30^. We employed HRMS in DIA mode that permitted stepwise acquisition for the detection of adducts and is useful for its archival nature as samples that have been analyzed earlier can be re-examined for new adducts. Even with the fragmentation data being limited to MS/MS, DIA approach has made significant progress in proteomics^31,32^ and metabolomics^33,34^. Development of bioinformatics tools related to adduct analysis would further enhance the use of this method in adductomics.

In the detection strategy, we utilized the common dR fragment present on nucleosides. This analytical approach did not allow us to capture rapidly depurinating adducts^35^ and phosphate-adducts formed on the phosphodiester backbone of DNA^36^. Further, bulky adducts such as those formed from covalent binding of nucleophilic sites on DNA to reactive metabolites of benzo[a]pyrene and other PAHs, having high molecular mass (□ 350 g/mol), were not targeted in this work. These adducts require metabolic biotransformation of the precursor PAH^37^, the ability of which remains to be explored for amphipods and other relevant species.

A relatively large amount of DNA from individual amphipods was obtained for this work; 30 to 105 µg/female and 15 to 95 µg/brood. We did not attempt linking the nucleoside adducts to a specific type of embryo malformation, because the DNA was extracted from whole broods, with 13 to 58 embryos, often with mixed malformation types. For establishing whether specific DNA modifications occur in an embryo with a certain malformation type, the samples have to be prepared using fewer embryos carrying the malformation type in question. This would require a higher sensitivity for detection of adducts and, for instance, could be done by using nano-LC flow condition utilizing the inverse relationship between flow rate and electrospray sensitivity^38^.

Amongst the 23 putative adducts detected, the chemical structure of 8 nucleoside modifications was proposed (**Fig. 2**) based on the elemental composition determined from the high mass accuracy, and the identity of 3 adducts was confirmed by comparison with commercially obtained standards. Significantly, two epigenetic-type changes in the female DNA, 5-me-dC (A3) and N^6^-me-dA (A22), were identified in this work as predictors for producing malformed embryos. DNA methylation is a characteristic feature for control of gene expression and maintaining genomic stability. This type of epigenetic alteration has been implicated in numerous biological processes, such as cell differentiation, X-chromosome inactivation, embryogenesis and tumorigenesis^8,39-41^. Many environmental contaminants in the Baltic sediments, including metals (e.g., Cd, Zn, and Hg), polychlorinated biphenyls and PAHs^16,17^, might interfere with DNA methylation status. Specifically, 5-me-dC has commonly been used as an epigenetic marker in evaluating the association of environmental chemicals to adverse health outcomes^6,42,43^. The N^6^-me-dA adduct is comparatively less studied, and its biological role is still unclear. However, the recent identification of N^6^-me-dA in frog *Xenopus laevis*, mouse *Mus musculus* and human tissues^44^, and in mouse embryonic stem cells^45^, suggest that this type of DNA methylation is more prevalent in the eukaryotes than previously thought. An unidentified DNA adduct (A9), having *m/z* 239.1139 of the nucleoside adduct was the most significant predictor of the health status. This mass does not seem to correspond to any methylated derivate of the nucleosides, suggesting another pathway contributing to the pathological development.

Also, three oxidative adducts were detected, 8-oxo-dG (A21), 5-OH-dC (A5) and 8-OH-dA (A8); they might be associated with the formation of ROS and cytochrome P450 isozymes. In particular, chromosomal aberrations and mutations, particularly GC to TA transversions, have been linked to the formation of 8-oxo-dG^5,46^. The oxidized guanine might also be associated with detrimental effects on cell function, such as microsatellite instability and acceleration of telomere shortening^4^. However, no significant correlation was observed between the 8-oxo-dG adduct and embryo damage.

We observed significant ontogenetic differences in the nucleoside adduct profiles. Comparison of the adduct profiles between the females and their embryos showed that some adducts (A15, A20, and A23) correlated significantly positively between the mothers and their offspring. However, no such correlations were observed for the adducts that were identified as significant predictors for the high frequency of the aberrant embryos (**Table S1**). The overall variability of the adduct profiles was significantly higher among the broods than among the females (**Fig. 4**), which was most likely related to both the frequency of the aberrations and the development-stage variability of the embryos. Although we targeted a relatively narrow span of the embryo development (stage 4–8) when selecting the gravid females for the adduct analysis, it is possible that organogenesis that occurs during the ontogenetic development during these stages was sufficient to induce the high variability in the relative distribution of the adducts in the embryos. It is also possible that embryonic cells are repair-efficient, and only a small number of DNA lesions remain unrepaired. As a result, none of those adducts that were detected in the embryos were retained in the predictive model that discriminated between the females carrying low and high proportions of the aberrant embryos.

The validation of DNA modifications for use as exposure biomarkers in environmental status assessment requires establishing causal relationships with pathological conditions, such as tumorigenesis or developmental aberrations. The latter was addressed in our study using field-collected animals, but more controlled experimental data relating profiles of DNA modifications, oxidative balance, and reproductive pathologies are needed. In the field, biological effects of contaminants typically result from exposure to complex mixtures of environmental contaminants with different bioavailability and a myriad of confounding factors, such as temperature, oxygenation, and feeding conditions. All these factors may affect rates of adduct formation and DNA repair. Standardizing exposure conditions would facilitate identification of unique, measurable nucleoside adducts indicative of exposure, elucidate which chemicals induce these adducts and provide information on the dose-response relationships. However, despite current knowledge gaps, DNA adductomics represents a very promising means for application in risk assessment and monitoring surveys. When more extensive data become available, they might be used to establish effective biological dose and the potential for irreversible toxicity, confirm suspected exposures and improve assessment of human impacts on the quality status of the marine environment.

## ACKNOWLEDGEMENTS

This work was financially supported by the Swedish Research Council Formas (2017-00864), the Erasmus+ program, the Department of Environmental Science and Analytical Chemistry at Stockholm University, and the United States National Institutes of Health (R01CA095039).

## AUTHORS CONTRIBUTIONS

HVM and EG conceptualized the project; BS performed the embryo analysis; NHM performed the DNA extraction; GM and HVM performed the adductomics; EG performed the statistical analysis; GM, HVM, and NYT conducted the adduct identification; HVM and EG wrote the manuscript with input from NYT. All authors reviewed the manuscript.

