## Supplementary information for "DNA epigenetic marks are linked to reproductive aberrations in amphipods"

**Figure S1. Map showing the locations in the Baltic region from where the amphipods are collected.** The stations marked in black are those monitored yearly within SNMMP and from where the amphipods were collected for this work, and those marked in red are stations selected for effect screening but not part of this study.

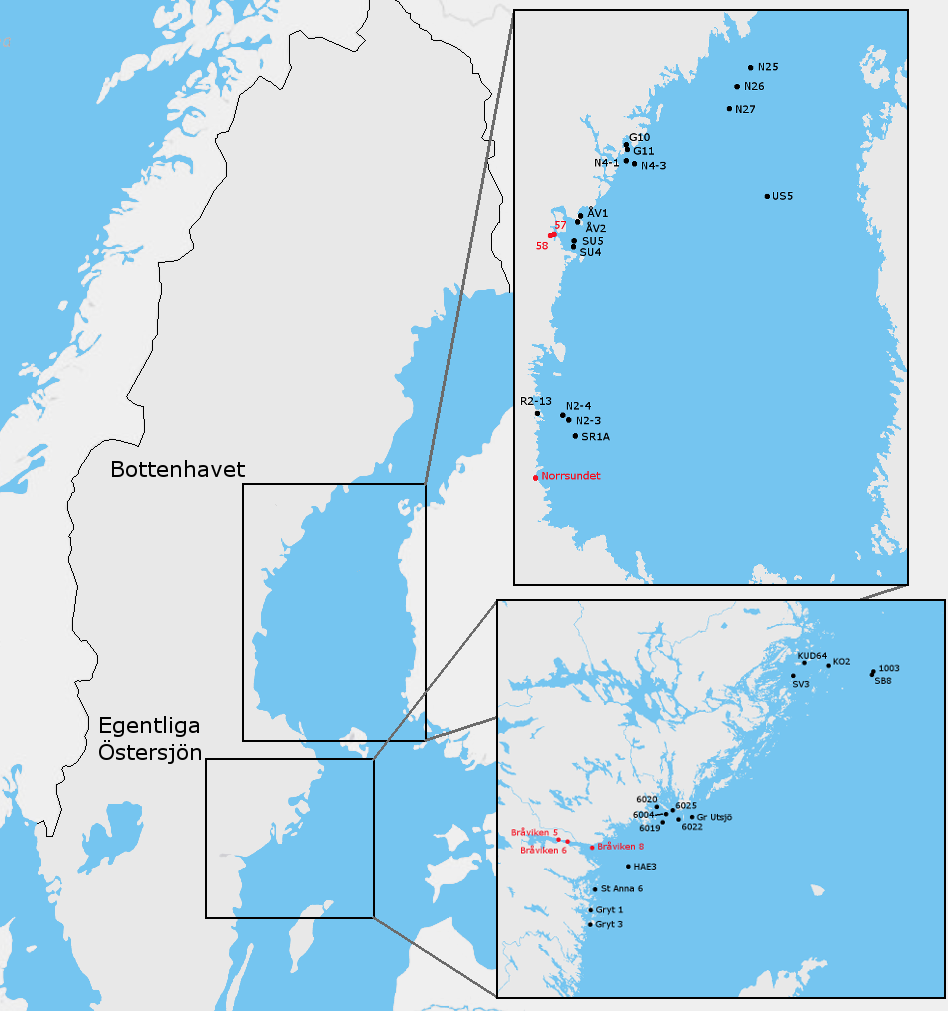

**Figure S2. Illustration of nucleoside adduct (M) and its fragmentation pattern.** The 2'-deoxyribose and the modification moieties are shown as dR and A, respectively. The protonated dR fragment, with m/z 117.0552, was used for screening of the adducts. The identification of adducts was based on the nucleobase fragment containing A as they are specific to the individual adducts.

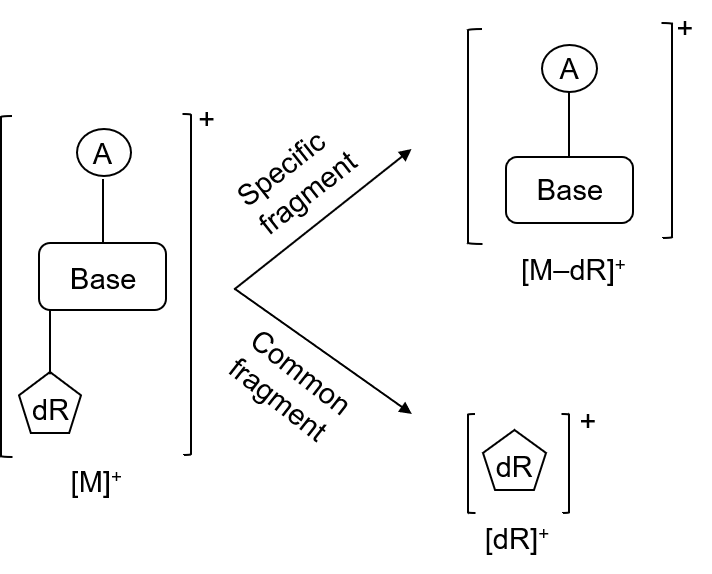

**Figure S3. Evaluation of the method used for quantification of nucleoside adducts.** ctDNA (n=5) was digested and analysed using the same method as that applied for the amphipod samples. The normalized mean values from the 5 samples are represented in the plot, and the standard deviations are shown as respective error bars. For the measured adducts in ctDNA, the method was shown to be repeatable with CV ˂10% indicating a low variability in the analysis.

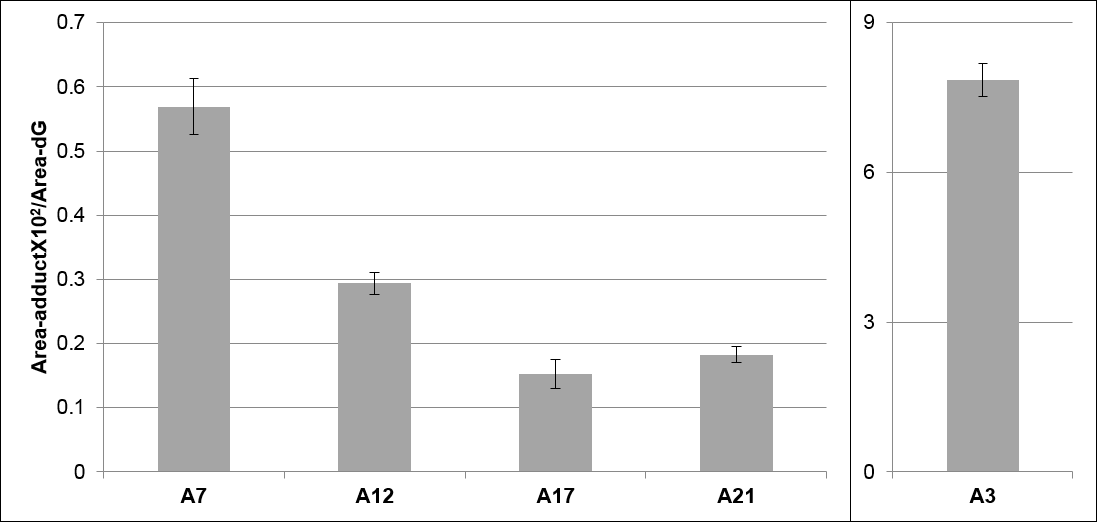

**Figure S4. Identification of 5-me-dC (A3), N^6^-me-dA (A22) and 8-oxo-dG (A21) in amphipods DNA using respective standards.** An overlap of peaks corresponding to the nucleoside adducts in amphipod samples before (continuous curve) and after (dashed curve) spiking of respective standards, using m/z of respective [M-dR+H]^+^, confirmed the identification. EIC for A21 in amphipod samples showed two peaks, but only the second peak had a retention time similar to that of the standard indicating the first peak to be an artefact.

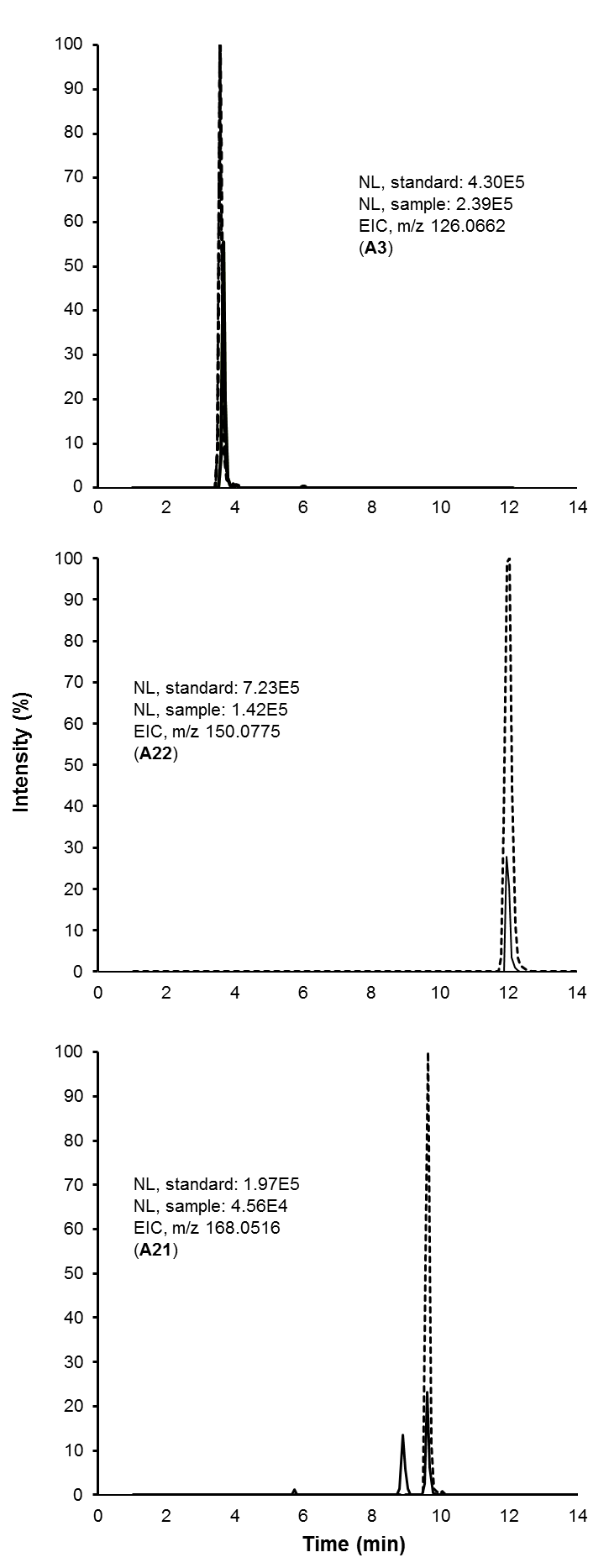

**Figure S5. Distribution of the measured nucleoside adducts (as well as dA, dC and T) shown as histograms and box-plots.** Scale of all adduct values correspond to peak areas normalized to dG (adduct area × 10^2^ / dG area) for (**A**) females and (**B**) embryos. The box represents 25%–75% of the values, with median shown as a horizontal line. The center of the rhombus represents the mean, and its upper and lower edges represent 95% confidence interval for the mean. The whiskers show non-outlier range, and outliers are displayed as dots. Adducts that were below the apparent quantification limit of the instrument (A5 and A10 in females; A1, A5, A10, and A22 in embryo broods) as well as those attributed as sodiated adducts (A4, A13, A16, and A18) and having high correlation with corresponding protonated species are not shown.

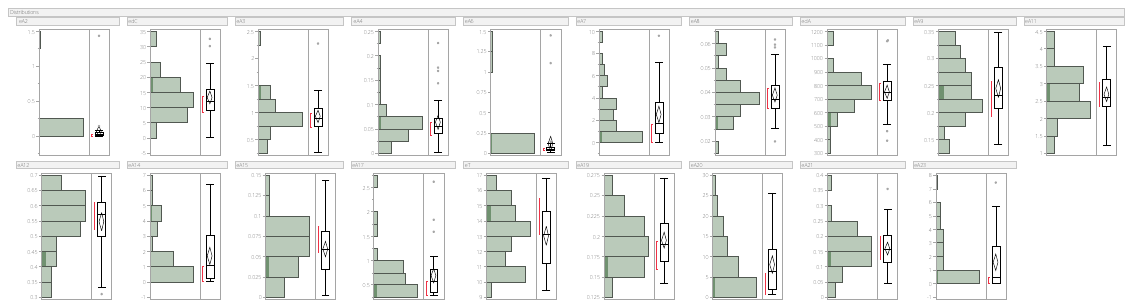

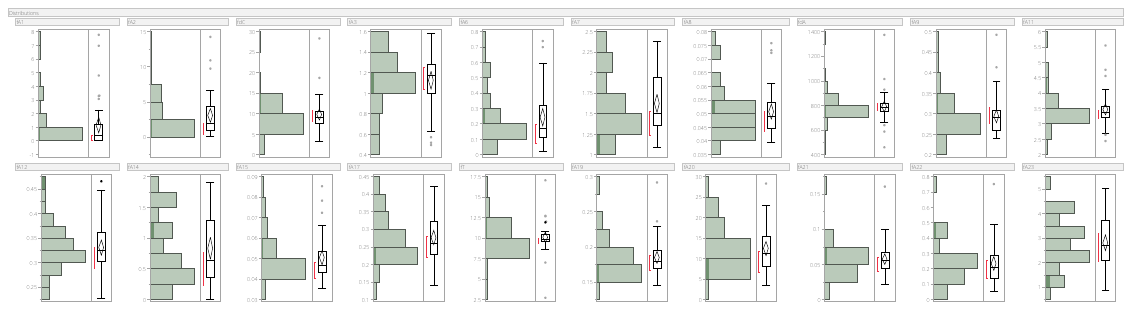

**B**

**A**

**Figure S6. Variations in the normalized peak area for nucleoside adducts (as well as dA, dC and T).** Measured in the *M. affinis* females and their embryos; in total, 40 gravid females were used generating the same number of samples for each group. Scale of all adduct values correspond to peak areas normalized to dG (adduct area × 10^2^ / dG area) and a two-way grouping is done, i.e. embryo (E) and female (F), and healthy (H, left, in blue) and unhealthy (U, right, in red). If >5% of embryos in the brood pouch were malformed, both the female and her embryos were classified as unhealthy, otherwise, as healthy; n = 19 and 21 for U and H, respectively. Box and whiskers indicate interquartile range and limits for the 95%-confidence interval, respectively; data points above or below the whiskers represent possible outliers. Adducts (F: A5 and A10; E: A1, A5, A10 and A22) that were below the apparent quantification limit of the instrument are not shown.

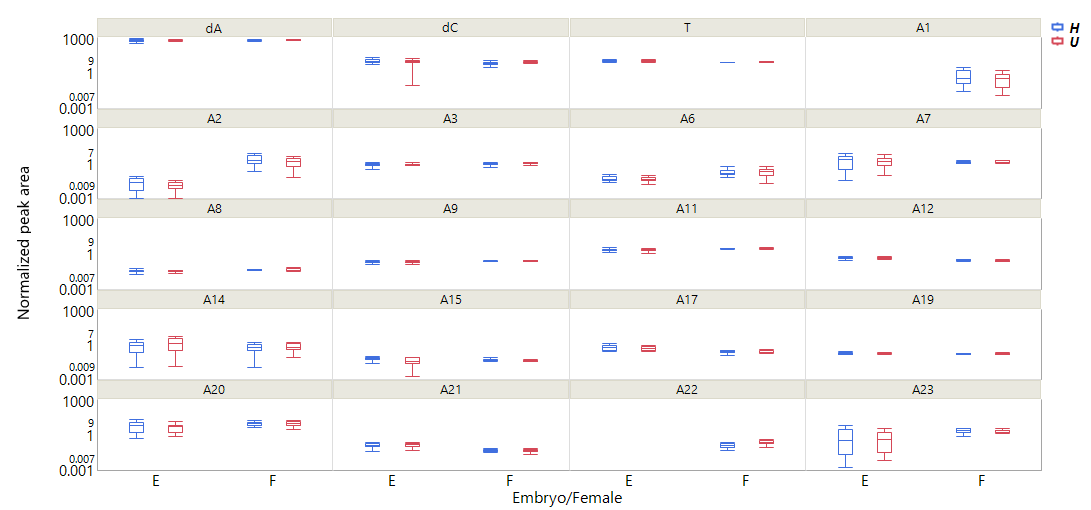

**Figure S7. Cross-correlation matrix for nucleoside adducts in the *M. affinis* using pooled data for the female and embryo samples.** Pearson *r* for Box-Cox transformed data for normalized peak area was used; those adducts that were below the apparent quantification limit were excluded.

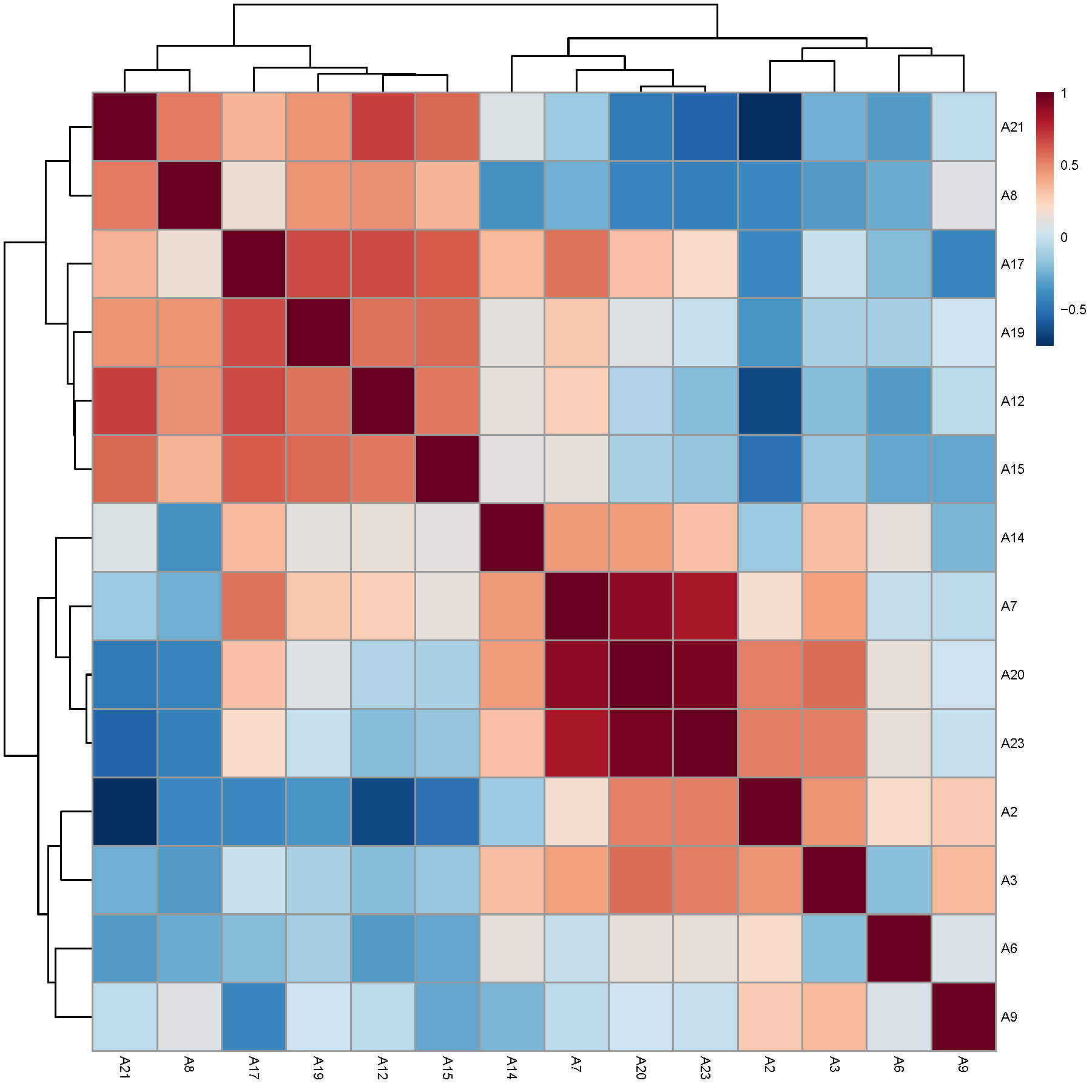

**Figure S8. Discrimination of the adductome in healthy and unhealthy amphipods based on area under the receiver operating characteristic (ROC) curve (AUC) logistic regression approach.** For each of the top three discriminating nucleoside adducts, the left panel shows the AUC confidence interval, true positive and false positive rates, and confidence interval (CI), the right panel shows the normalized values for the adducts in healthy and unhealthy individuals. AUC logistic regression approach identified female A22 (AUC = 0.802), A9 (AUC = 0.697), and A3 (AUC = 0.652) to have the greatest specificity and sensitivity for distinguishing the adductome of reproductive pathologies in the amphipods.

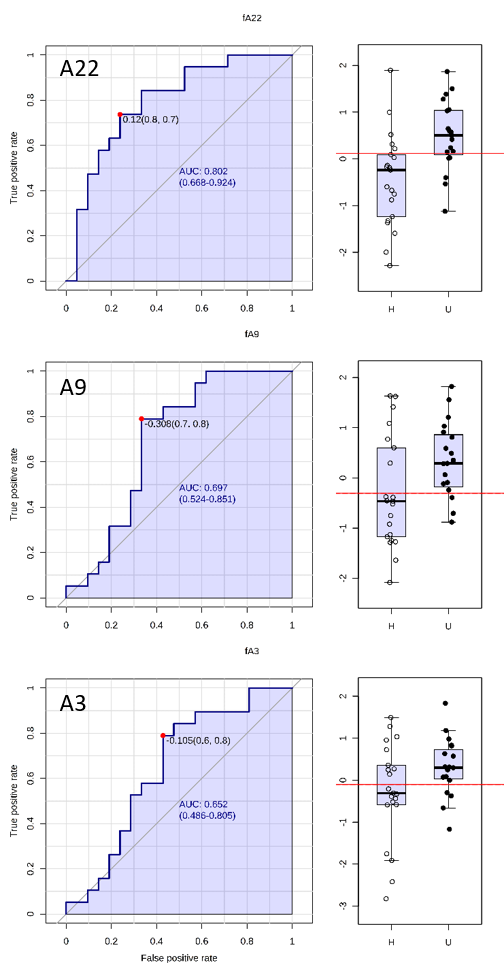

**Table S1. Cross-correlation (Pearson r) for specific nucleoside adducts between the female and the embryo DNA.** The adducts (A1 to A23) were measured in the females (rows: fA1 to fA23) and their embryos (columns: eA2 to eA23); altogether, 40 paired female-brood samples were used. The data were Box-Cox transformed; the significant (p < 0.05) correlations are in bold. Significant female-embryo correlations for specific adducts (A15, A20 and A23) are shown in red.

|  | **eA2** | **eA3** | **eA6** | **eA7** | **eA8** | **eA9** | **eA11** | **eA12** | **eA14** | **eA15** | **eA17** | **eA19** | **eA20** | **eA21** | **eA23** |
| --- | --- | --- | --- | --- | --- | --- | --- | --- | --- | --- | --- | --- | --- | --- | --- |
| **fA1** | 0.09 | 0.24 | -0.26 | 0.13 | -0.02 | 0.24 | 0.21 | 0.26 | -0.14 | 0.18 | 0.05 | 0.01 | 0.00 | 0.23 | 0.07 |
| **fA2** | 0.04 | 0.18 | -0.29 | -0.11 | 0.06 | **0.44** | **0.41** | **0.45** | **-0.47** | -0.07 | -0.14 | -0.11 | -0.11 | 0.30 | -0.29 |
| **fA3** | 0.06 | -0.11 | 0.30 | **-0.31** | -0.07 | -0.25 | -0.13 | -0.16 | -0.01 | **-0.31** | -0.15 | -0.00 | -0.18 | -0.07 | -0.31 |
| **fA6** | -0.08 | -0.06 | 0.14 | **0.36** | -0.03 | -0.22 | 0.02 | -0.19 | **0.49** | 0.25 | **0.32** | 0.27 | 0.28 | 0.01 | **0.47** |
| **fA7** | 0.08 | 0.01 | -0.04 | 0.22 | 0.23 | -0.19 | -0.09 | -0.08 | 0.04 | 0.16 | 0.19 | -0.10 | 0.23 | -0.06 | 0.30 |
| **fA8** | 0.05 | 0.01 | -0.17 | -0.08 | 0.07 | -0.12 | 0.04 | 0.02 | -0.22 | -0.06 | -0.03 | -0.16 | 0.04 | 0.07 | -0.03 |
| **fA9** | -0.08 | -0.06 | 0.02 | -0.21 | 0.00 | -0.05 | 0.11 | 0.08 | 0.08 | -0.21 | -0.09 | 0.09 | -0.17 | 0.11 | -0.24 |
| **fA11** | -0.00 | -0.00 | -0.08 | -0.09 | 0.09 | -0.09 | -0.04 | 0.06 | -0.01 | -0.18 | -0.07 | -0.06 | -0.07 | -0.07 | -0.04 |
| **fA12** | 0.12 | -0.01 | -0.19 | 0.10 | 0.13 | -0.22 | -0.18 | -0.10 | -0.01 | 0.01 | 0.14 | -0.07 | 0.19 | -0.16 | 0.16 |
| **fA14** | -0.03 | -0.19 | 0.09 | -0.26 | 0.05 | -0.10 | 0.06 | -0.03 | 0.05 | -0.28 | -0.15 | -0.05 | -0.04 | -0.20 | -0.23 |
| **fA15** | 0.21 | 0.09 | -0.14 | **0.32** | -0.10 | -0.19 | -0.19 | -0.21 | 0.09 | **0.35** | 0.22 | 0.02 | 0.22 | -0.00 | **0.34** |
| **fA17** | 0.02 | 0.07 | -0.20 | 0.17 | -0.08 | 0.02 | 0.06 | 0.02 | -0.18 | 0.04 | -0.08 | -0.30 | 0.13 | 0.15 | 0.09 |
| **fA19** | -0.01 | 0.03 | -0.17 | 0.08 | -0.06 | 0.01 | 0.06 | 0.03 | -0.13 | -0.07 | -0.06 | -0.09 | 0.04 | 0.22 | 0.02 |
| **fA20** | 0.15 | 0.03 | 0.03 | 0.31 | 0.30 | -0.30 | -0.15 | -0.14 | 0.13 | 0.22 | **0.32** | 0.03 | **0.36** | -0.14 | **0.42** |
| **fA21** | 0.12 | -0.01 | -0.02 | **0.41** | 0.03 | **-0.38** | -0.15 | -0.29 | **0.44** | 0.17 | **0.32** | 0.18 | **0.39** | -0.05 | **0.43** |
| **fA22** | -0.04 | -0.30 | 0.26 | -0.11 | 0.02 | -0.23 | -0.13 | -0.04 | 0.08 | -0.25 | 0.00 | 0.04 | 0.07 | -0.15 | -0.11 |
| **fA23** | 0.10 | 0.13 | -0.12 | 0.45 | 0.07 | -0.05 | -0.05 | 0.07 | 0.05 | 0.28 | 0.24 | -0.10 | 0.27 | -0.07 | **0.52** |

**Table S2. Logistic regression output.** Classification of cases for unhealthy (U) and healthy (H) females. Odds ratio: 22.666667, Log odds ratio: 3.120895.

| Observed | Predicted - U | Predicted - H | Correct, % |
| --- | --- | --- | --- |
| U, 19 | 16 | 3 | 84.2 |
| H, 21 | 4 | 17 | 80.9 |
